# MetaboMiNR: a web application to analyze label free quantitative proteomic experiments with a focus on metabolism and nuclear receptors

**DOI:** 10.1101/2025.03.25.644590

**Authors:** Michael F. Saikali, Carolyn L. Cummins

## Abstract

Label free quantitative (LFQ) proteomics is growing in popularity and becoming increasingly more accessible to researchers, empowering them to compare proteome-wide changes between different treatment conditions. However, it remains difficult to leverage the full potential of LFQ data when the researcher has limited experience in proteomics and/or bioinformatics due to the complexity of data analysis. Here, we present MetaboMiNR, an easy-to-use web application for the analysis of LFQ data with a focus on metabolism and nuclear receptors. MetaboMiNR guides users through an intuitive process with clear instructions and minimal user input to conduct statistical analysis and produce publication ready plots. Users may input a MaxQuant generated output file and perform standard global analysis and data quality control with the click of a button. The application offers 3 additional unique features: 1) Metabolism Miner extracts a user selected Reactome pathway from the dataset, 2) Nuclear Receptor Miner extracts the target genes of a user-selected nuclear receptor, and 3) Individual Plotter produces publication-ready bar plots for a selected protein. The utility of this application was demonstrated by analyzing a previously published dataset from mice treated with LDT409, a synthetic PPAR agonist. MetaboMiNR can be accessed freely at https://cumminslab.shinyapps.io/MetaboMiNR/.

---

1. **Introduction**

Nuclear receptors (NRs) are a family of 48 transcription factors in humans that have been extensively studied because of their critical roles in endocrine signaling^1^. NRs are activated by small-molecule ligands and control the expression of downstream target genes involved in myriad processes including development, reproduction and metabolism^1,2^.

In 2002, the Nuclear Receptor Signalling Atlas (NURSA)^3^ was formed to collate and share large datasets between investigators who were generating genome-wide proteomic and transcriptomic data related to NRs. These NR specific datasets have since been incorporated into the expanded Signalling Pathways Project (SPP)^4^ knowledgebase that creates tools to allow bench researchers to mine integrated transcriptomic and cistromic data. Of relevance to NR researchers is the ‘consensome’ tool that summarizes the community’s consensus for a NR reference gene signature^4,5^. For scientists studying NRs, RNA-seq and shotgun proteomics is becoming an increasingly accessible tool to phenotype and understand NR signal transduction. Our research group and countless others have used a combination of these tools to develop a clearer understanding of NR signalling from the activation of gene transcription to protein production and functional outcomes^6–9^.

Mass spectrometry (MS) based bottom-up proteomics is an increasingly popular laboratory technique to identify changes in the proteome between test conditions^10,11^. In a typical bottom-up proteomics experiment, proteins from a biological sample are digested with a protease that will produce peptides that can be predicted *in silico* from the defined protease cleavage site and sample species proteome^12,13^. These digested peptides are chromatographically separated by nanoflow liquid chromatography (nLC) and detected by a high-resolution MS to produce a unique mass spectrum for each peptide sequence^14,15^. Software such as MaxQuant will match these spectra to peptide sequences which will be mapped back to protein sequences and combine their intensities using algorithms developed for label-free quantification (LFQ)^16–18^. This technique allows for the identification and quantification of proteins. When combined with upstream experiments such as drug treatment or genetic deletion it can be used to determine the effect these interventions had to the proteome^19^.

Beyond sample processing and instrument time, data analysis is a significant bottleneck in the ’omic pipeline and can be daunting for labs that are new to proteomics. While downstream analyses of the data may differ between experiments, many experiments require the same global unbiased analysis up front. There is no shortage of software options for proteomic analyses, with many being free (e.g., Perseus^20^). Software such as Perseus generally include a vast toolbox for analysis from the simplest t-test to complex visualizations. However, as core facilities help make proteomics more commonplace, these vast toolboxes can leave an inexperienced end user feeling overwhelmed.

In this paper we present MetabomiNR, a web application designed to analyze LFQ proteomic data and generate publication ready plots with a focus on NR target genes and metabolism. The application conducts standard global proteomic analyses, identifies differentially expressed proteins, while also mining the dataset for genes related to pathways in intermediary metabolism, and NR target genes. To demonstrate the utility of this application, we apply it to a previously published dataset from our group in which we investigated the impact of LDT409 treatment, a novel pan agonist of the peroxisome proliferator-activated receptors (PPARs), on mouse liver protein expression.

## 2. Methods

### 2.1 Global Analysis Pipeline

To perform the initial global proteomic analysis, MetaboMiNR has leveraged the DEP application package in R that was developed for Differential Enrichment analysis of Proteomics data^21^. Data are imported using native DEP functions and a downloadable template is then generated containing the sample names along with columns for the condition, replicate number, and order number. The condition file is then completed by the user and uploaded back into the application and analysis may begin. Native DEP functions are then used to identify missing values and impute them, with the application allowing for user input to control the threshold setting for missed values. The default threshold is 0 which means that for a protein to be kept, it must be detected in all replicates of at least one condition. Increasing the threshold decreases that requirement by 1. For example, in a 4-replicate experiment if the threshold is set to 1, it will require the protein to be detected in at least 3 out of 4 of the replicates, in at least one condition.

MetaboMiNR conducts statistical analyses using the DEP package, while presenting the user with an interface allowing for input of parameters such as the *P*-value cut off, log_2_FC cut off, along with the condition on the right (numerator), and the condition on the left (denominator) (refer to item 10 in the Supplemental User Guide for further details). The application presents a PCA analysis, which with DEP defaults to the top 500 most variable proteins. A sliding scale was added so that the user can change the number of proteins included in the PCA analysis. The upper limit is set to the total number of proteins detected in that dataset. Finally, this global analysis pipeline produces a heatmap of the differentially expressed proteins while allowing the user to customize the scale of the z-score and the number of defined clusters.

### 2.2 Metabolism Miner

The output from the global analysis pipeline is continued onto the Metabolism Miner tab. The application accesses the gene lists found in Reactome pathways^22^ and extracts those genes out of the data set, allowing the user to select specific pathways to investigate. Reactome by default exports human genes, so conversion to their *mus musculus* orthologues is required when analyzing datasets that were performed in mice. By default, the user is provided a curated list of metabolic pathways for easy access, but the ReactomeID for any pathway can be submitted for automated gene list extraction and analysis. The data are presented as a heatmap and are also downloadable as a tab-delimited file in case the user wants to perform additional offline analyses.

### 2.3 Nuclear Receptor Miner

Similar to the Metabolism Miner, the Nuclear Receptor Miner uses the concensomes provided by the SPP^4^ to extract likely target genes of the selected NR. These concensomes are generated from compiled cistromic and transcriptomic data to produce a list of likely target genes for the receptor of interest. The user can select a receptor and apply a percentile cut-off to control the size of the gene list to be extracted. A higher percentile cut-off (>95) will ensure that only high-confidence NR target genes will be included. If these gene names are present in the protein dataset, they will be presented as part of a heatmap.

### 2.4 Individual Plotter

The Individual Plotter tool allows the user to search the dataset for a specific gene either with the UniprotID or the gene symbol. The data are presented to the user as a bar plot with standard error. The user is able to adjust the order and colors of the bars. While the bar plots do not directly display adjusted *P*-values, the data table under the plot contains all the *P*-values for all performed comparisons.

### 2.5 Proteomics & Data Availability

A published dataset from our group was used as a test case. Full details of the dataset may be found in the previously published article^7^. Briefly, mouse liver samples were collected from 6-month-old C57BL/6 mice fed a normal chow diet or a high fat high cholesterol diet (HFD, Envigo TD88137) for 6.5 weeks. During the last 18 days the mice were intraperitoneally injected with vehicle or 40 mg/kg LDT409 a synthetic PPAR pan agonist. Samples were processed with the filter aided sample preparation protocol^23^ and desalted on in-house made C18 pipette tips. Analysis was done on an EASY nLC1200 coupled to a Thermo QExactive HF run in top20 data dependent acquisition (DDA) mode. The data were processed through MaxQuant v1.6.10.43 and can be accessed from the PRIDE repository with the identifier PXD047662.

## 3. Results

### 3.1 Global Analysis of LDT409’s effect on chow and HFD fed mouse liver proteomes

The MaxQuant output of the LDT409 dataset was uploaded to MetaboMiNR and processed using the condition file in **Table 1**. The dataset was filtered for proteins that were detected in all replicates of at least one condition (**Figure 1A,B**) resulting in 1918 proteins. The missing values were imputed, and an FDR corrected t-test was conducted between every condition to determine the differentially expressed proteins (DEPs). In total, 167 DEPs were identified as change in at least one condition. MetaboMiNR automatically conducts a principal component analysis (PCA) on the top 500 most differentially expressed proteins. This analysis revealed clustering of the biological replicates and distinct separation of the conditions into 3 groups (**Figure 1C**). It showed a clear difference in the liver proteome between high fat diet and normal chow fed mice along PC1 while LDT409 treatment formed its own group along PC2 with mild separation between the diets.

**Figure 1:**
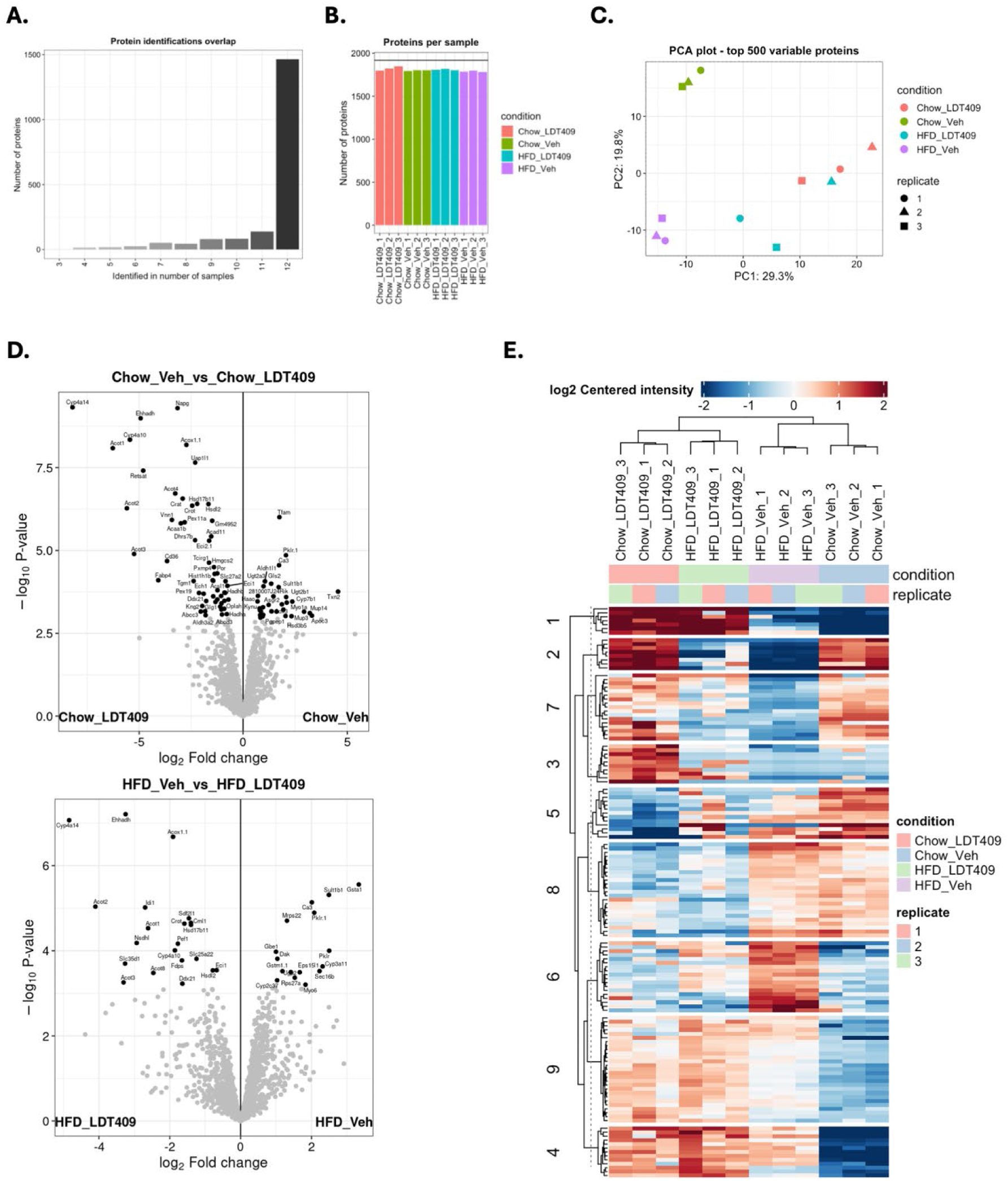
Global Analysis reveals LDT409 and HFD-specific effects on the mouse liver proteome. Global analysis of the LDT409 dataset. **A.** Summary of protein identification overlap with a threshold set to 0 meaning that for a protein to be kept, it must be detected in at least every replicate of a single treatment condition. **B.** Summary of proteins identified per sample. **C.** Principal component analysis (PCA) on the top 500 differentially expressed proteins. **D.** Volcano plot comparing the Chow Veh group the Chow LDT409 group (top) and HFD Veh group to the HFD LDT409 group (bottom). Points that meet the criteria of log_2_FC > 0.5 and *P*_adj_ < 0.05 are highlighted and labeled in black. **F.** Heatmap of the differentially expressed proteins clustered into 9 groups.

**Table 1:** Completed conditions template. This table was produced by MetaboMiNR after uploading the LDT409 proteinGroups.txt file. The condition and replicate columns were filled to indicate the sample groups and replicate numbers.

| label | condition | replicate |
| --- | --- | --- |
| MS01 | Chow_Veh | 1 |
| MS02 | Chow_Veh | 2 |
| MS03 | Chow_LDT409 | 1 |
| MS04 | Chow_LDT409 | 2 |
| MS05 | HFD_Veh | 1 |
| MS06 | HFD_Veh | 2 |
| MS07 | HFD_LDT409 | 1 |
| MS08 | HFD_LDT409 | 2 |
| MS09 | HFD_Veh | 3 |
| MS10 | HFD_LDT409 | 3 |
| MS11 | Chow_Veh | 3 |
| MS12 | Chow_LDT409 | 3 |

MetaboMiNR presents the user with a visualization for the statistical analysis in which a comparison can be selected, and it will be automatically plotted as a publication ready volcano plot. This feature was used to identify LDT409 induced DEPs in both diets (**Figure 1D**). Hepatic CYP4A14, EHHADH, ACOT1, ACOT2, ACOT3, CYP4A10 and 7 additional proteins were all significantly upregulated upon LDT409 treatment in both high-fat and chow fed mice. DAK, CYP2C37, SULT1B1, CA3, and PKLR.1 were the only proteins consistently supressed by LDT409 treatment in both diets. The final step of the global analysis is a k-means clustered heatmap of DEPs allowing the user to identify meaningful expression patterns. LDT409 was shown to reverse HFD-induced metabolic associated steatotic liver disease, obesity, and insulin intolerance^7^. To identify proteins that may play a role in this correction, we used the k-means clustered heatmap to identify proteins that are dysregulated in the HFD control group but rescued by treatment with LDT409 (**Figure 1E**). Cluster 2 represents proteins that are repressed by HFD but moderately rescued by LDT409 treatment. Similarly, cluster 6 represents proteins where the HFD induction is lost with LDT409 treatment. Clusters 1 and 9 contain proteins that are induced by LDT409 irrespective of diet. In summary, the global analysis page was used with minimal user input to process the data, identify DEPs, and identify meaningful expression patterns specific to the experimental question.

### 3.2 The Metabolism Miner function reveals LDT409 as an activator of mitochondrial β-oxidation and peroxisomal lipid metabolism

The next tool in MetaboMiNR is the Metabolism Miner. This feature allows the user to select from a curated list of Reactome pathways^22^ or enter the reactomeID of any pathway and automatically extract from the data proteins within the selected pathway. Considering that LDT409 had been identified as a pan PPAR agonist^24^, we used MetaboMiNR to search for proteins involved in the synthesis of ketone bodies (R-HSA-77111), mitochondrial β-oxidation (R-HSA-77289), and peroxisomal lipid metabolism (R-HSA-390918) along with cholesterol biosynthesis (R-HSA-191273) which is expected to change with diet but not PPAR activation.

MetaboMiNR extracted the proteins involved in each of these Reactome-defined pathways and displayed the results as a heatmap (**Figure 2**). Out of the 8 proteins in the synthesis of ketone bodies term, 6 were found in the dataset and did not display an obvious treatment effect (**Figure 2A**). The cholesterol biosynthesis term contained 14 of the 25 proteins in the pathway and displayed a striking HFD-induced repression (**Figure 2B**). The mitochondrial β-oxidation pathway exhibited an LDT409 effect divided into two clusters; the top cluster representing proteins induced by LDT409, and the bottom cluster representing proteins induced by HFD and rescued by LDT409 treatment (**Figure 2C**). Finally, peroxisomal lipid metabolism pathway exhibited a treatment effect on many of the extracted proteins.

**Figure 2:**
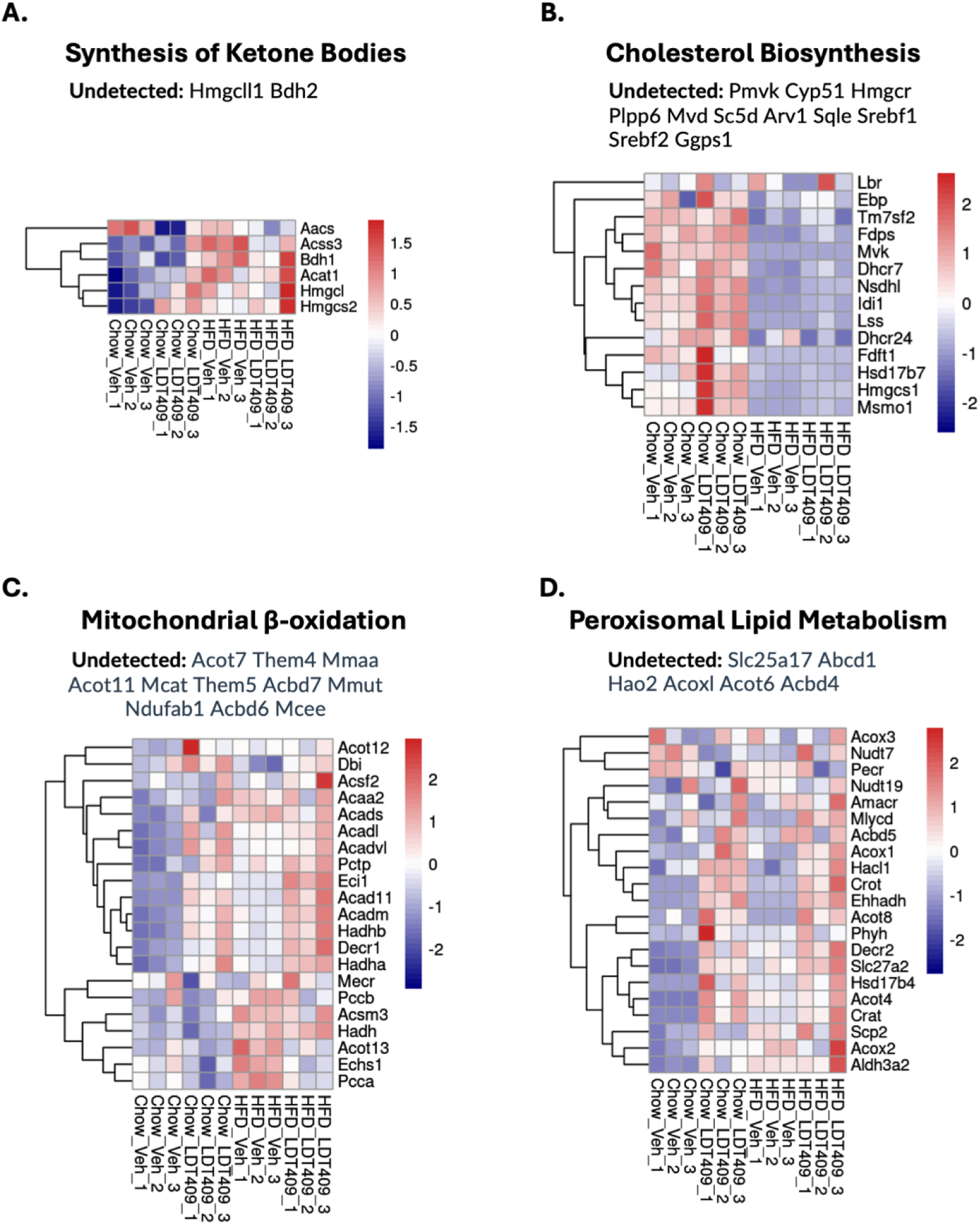
Metabolism Miner reveals LDT409 activates peroxisomal lipid metabolism in both chow and high-fat diet fed mice. Metabolism Miner was used to extract the proteins from the following pathways: **A.** Synthesis of ketone bodies (ReactomeID: R-HSA-77111) **B.** Cholesterol biosynthesis (ReactomeID: R-HSA-191273) **C.** Mitochondrial β-oxidation (ReactomeID: R-HSA-77289) **D.** Peroxisomal lipid metabolism (ReactomeID: R-HSA-390918). MetaboMiNR displays all of the genes in the selected pathway above the heatmap. This figure was edited to only report the proteins that were not detected in the dataset. The heatmaps were produced directly from MetaboMiNR.

These findings demonstrate the power of MetaboMiNR at allowing the user to rapidly extract proteins from their dataset allowing for fine-tuned interrogation of the results based on the hypotheses being tested or the observed phenotypes in the experiment.

### 3.3 The Nuclear Receptor Miner function identified PPAR as the most active NR upon LDT409 treatment among the available NR consensomes

MetaboMiNR uses the consensomes from SPP to create lists of potential target genes for the available NRs. At the time of publication these were AR, ER, ERR, FXR, GR, MR, PPAR, PR, RAR, ROR, THR, TR, and CAR/PXR. The default threshold is to select the genes in the top 95^th^ percentile of the consensome. We selected the PPAR consensome and MetaboMiNR automatically produced a heatmap of proteins that were detected and are flagged as PPAR target genes (**Figure 3A**). At a glance, it appears that LDT409 does exhibit a PPAR effect when compared to the vehicle and LDT409 treatment groups. The raw data can be downloaded for downstream experiment-specific analyses.

**Figure 3:**
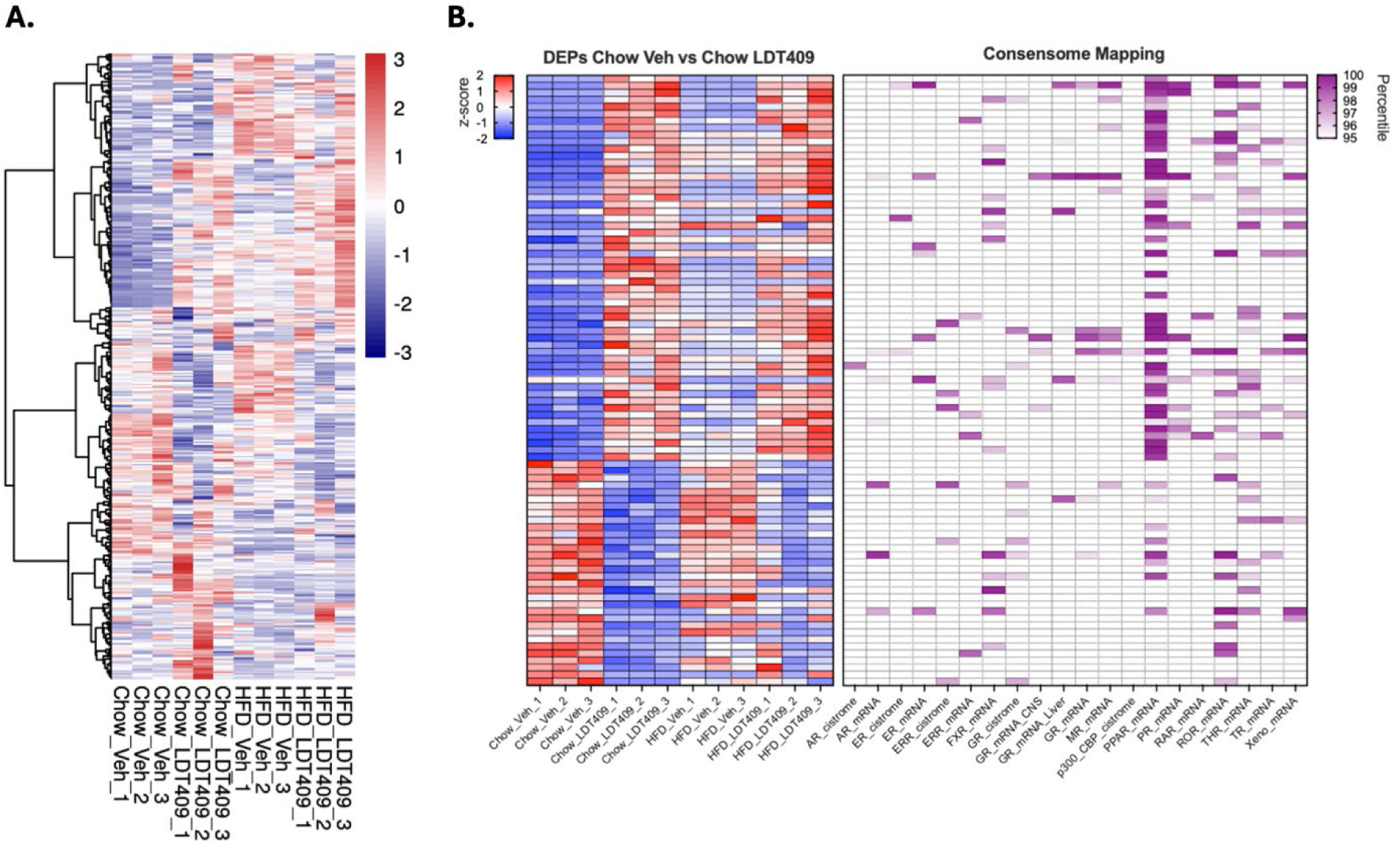
Nuclear Receptor Miner reveals PPAR as the most activated NR consensome by LDT409. **A.** Nuclear Receptor Miner extracted the top 95^th^ percentile of PPAR consensome genes from the LDT409 dataset and plotted the results as a z-scored heatmap. **B.** The Nuclear Receptor Miner results were downloaded and filtered for proteins that were induced by LDT409 in chow fed mice (*P_adj_* <0.05). The results were plotted as a z-scored heatmap on the left while a second heatmap was made with the consensome percentile scores. The cells shaded in white did not meet the 95^th^ percentile threshold, while those in purple do. Both heatmaps in **B.** were generated outside of MetaboMiNR.

To investigate if LDT409 is resulting in a change in the proteome of PPAR target genes, we downloaded the consensome-matched data from MetaboMiNR (**Supplemental File 1**) and filtered the results for DEPs between chow vehicle and chow LDT409. We then generated a heatmap of these results along with their consensome annotations (**Figure 3B**). LDT409 induced 55 proteins, 44 (74%) of which were identified as PPAR targets by MetaboMiNR based on their consensome score and the applied threshold. LDT409 repressed 32 proteins, with 9 (28%) representing PPAR targets. In total, there are 399 PPAR target genes within the 1918 proteins detected suggesting that 20.8% of a random selection of proteins should be PPAR targets. Comparing this base rate to the rate of observed target genes (74% and 28%), we can conclude LDT409 induces the protein expression of PPAR target genes.

### 3.4 The Individual Plotter allows for rapid production of publication ready plots

The final feature of MetaboMiNR is the Individual Plotter. It was designed with rapid re-analysis in mind allowing investigators to search for an individual protein from the dataset on command. Used in tandem with the Metabolism and NR Miner features, the pathways described above can be further investigated by plotting individual proteins as a bar graph.

3-hydroxy-3-methylglutaryl-CoA synthase 2 (HMGCS2) is the rate limiting enzyme in ketogenesis and *Hmgcs2* is a PPARα target gene in the liver. LDT409 induced the expression of HMGCS2 in the livers from chow fed mice but this increase was equally observed in response to high fat diet feeding and sustained with LDT409 treatment (**Figure 4A**). 3-hydroxy-3-methylglutaryl-CoA synthase 1 (HMGCS1) catalyzes the rate limiting step of cholesterol biosynthesis and was induced by LDT409 in chow fed mice (**Figure 4B**). HMGCS1 expression was significantly reduced by HFD feeding and was not recovered with LDT409 treatment. Acyl-CoA thioesterase 1 (ACOT1) is part of the mitochondrial β-oxidation pathway and was significantly induced by LDT409 regardless of diet (**Figure 4C**) as was EHHADH, a protein involved in peroxisomal lipid metabolism (**Figure 4D**).

**Figure 4:**
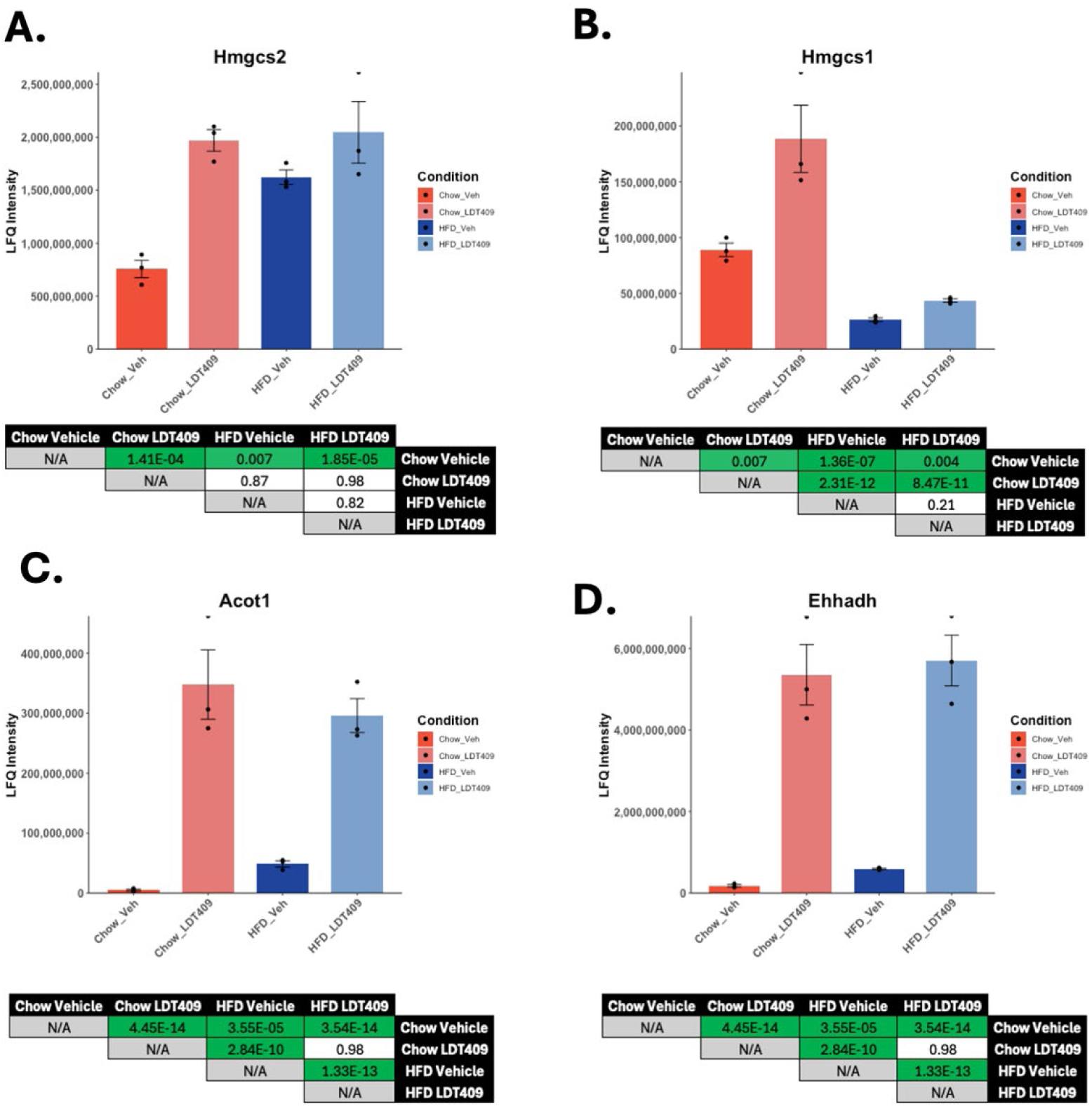
The Individual Plotter tool shows LDT409 induces the protein expression of many PPAR target genes. The Individual Plotter tool was used to plot a protein from each category in Figure 2; **A.** HMGCS2 (Synthesis of ketone bodies) **B.** HMGCS1 (Cholesterol biosynthesis) **C.** ACOT1 (Mitochondrial β-oxidation) **D.** EHHADH (Peroxisomal lipid metabolism). The bar plots were taken directly from the MetaboMiNR application. The *P_adj_* values listed for each comparison were obtained from the output table in MetaboMiNR and added to this figure outside of the program for easier interpretation.

## 4. Discussion

As proteomics becomes increasingly more accessible to research groups, applications for streamlining the analysis pipeline have been developed such as LFQ-Analyst^25^. These applications, including MetaboMiNR, are immensely useful for conducting a basic guided analysis of LFQ proteomics data. However, MetaboMiNR provides the additional unique advantage of interrogating the dataset with predefined pathway terms from Reactome or mining the data for NR target genes using the SPP consensomes. MetaboMiNR offers users a free, easy to use, streamlined and rapid analysis of LFQ proteomic experiments. The user only inputs the MaxQuant results file (proteinGroups.txt) and a conditions file for a which a template can be downloaded after uploading the MaxQuant data. Using the DEP package^21^, MetaboMiNR provides the user with a generalized global analysis in which the statistical results can be downloaded for further downstream specialized analyses. Furthermore, it generates publication ready volcano plots, heatmaps, and PCA plots. Any further user input is guided with on screen instructions designed to be intuitive and easy to follow without requiring background knowledge in proteomics. In this paper, we demonstrated both the general utility of this application and its utility for performing downstream analyses of metabolic pathways and NR-regulated pathways using our previously published LDT409 dataset.

LDT409, a partial pan-PPAR agonist, is a phenolic lipid derived from cashew nut shell liquid that is being investigated for its potential to treat diabetes and obesity^24^. Treatment with LDT409 has been shown to reduce diet-induced obesity, metabolic associated steatotic liver disease and insulin intolerance in both high-fat diet^7^ and high-fructose diet fed mice^26^. Using the global analysis tool, we were able to determine the general changes to the mouse liver proteome upon treatment with LDT409. The PCA plot revealed the separation of the chow and HFD vehicle groups from each other and from the LDT409 treated conditions. This is consistent with the observed phenotype that LDT409 rescues the liver from the effects of a high-fat diet. The proteome of the HFD LDT409 mice was more similar to the Chow LDT409 mice while remaining dissimilar from the Chow vehicle group suggesting LDT409 induces protective changes under a high-fat diet.

Using the statistical comparisons, we identified a set of 13 proteins that were consistently induced by LDT409 in both the HFD and chow fed mice (including CYP4A14 and CYP4A10). At the mRNA level, both genes have been demonstrated to be induced by PPARα through fasting, HFD feeding, and agonist activation^27^. Induction of CYP4A14 and CYP4A10 is greater after agonist treatment when compared to HFD feeding which is consistent with our observations at the protein level. ACOT1 was also consistently found to be upregulated by LDT409 and has been shown to be regulated by PPARα and HNF4α in the liver^28^.

The positive metabolic effects of PPARα agonism can be partly explained by their increased fatty acid uptake and oxidation in the liver^29–32^. The Metabolism Miner tool was used to identify the proteins in the dataset that are involved in both mitochondrial and peroxisomal β-oxidation. LDT409 induced the expression of ACADM, ACADVL, ACAD11, ECI1, HADHA, HADHB, ACOX1, and EHHADH, key enzymes in both mitochondrial and peroxisomal β-oxidation. The ability to easily visualize user-inputted Reactome pathways using MetaboMiNR will allow researchers more quickly assess the impact of their treatment on a signaling pathway and promote hypothesis generation.

Finally, the tool that truly distinguishes MetaboMiNR from other software is the ability to mine the dataset for nuclear receptor target genes. The Nuclear Receptor Miner tool matches the proteins detected in the dataset to the available NR consensomes producing a downloadable table containing the group comparison statistics, the normalized intensities, and a column for each NR consensome containing the percentile score. While the heatmap produced in the application is useful for a broad overview, the downloadable file is incredibly versatile as it allows the user to answer specific questions from their dataset with all of the statistical analyses already incorporated. We used this tool to investigate which NR consensome was most active upon LDT409 treatment. Consistent with its known PPAR activity, LDT409 appears to induce the protein expression of more PPAR target genes than any other NR. This analysis was done using the top 95^th^ percentile of each consensome which contained a total of 1203 PPAR target genes. The CAR/PXR and GR consensomes have a similar number of genes (1211 and 1165 respectively) but did not appear to be widely activated upon LDT409 treatment (**Figure 3B**) demonstrating LDT409’s specificity.

MetaboMiNR will be maintained as a free to use application updated annually. The Reactome pathways are requested live and will automatically update with Reactome, however the SPP consensomes will be periodically manually updated and expanded to include additional pathways. Overall, we demonstrated the utility and validated the results from MetaboMiNR using our LDT409 dataset. MetaboMiNR is a powerful tool that allows scientists interested in metabolism and NRs (with little to no experience in proteomics and bioinformatics) to produce meaningful analyses from their LFQ experiments.

## Supporting information

Supplemental User Guide

## Acknowledgements

We gratefully acknowledge Cigdem Sahin for preparing the LDT409 liver proteome samples, Adib Saikali for technical consulting, and the CLC lab members for testing the software and advising on user experience. Support for the project was provided by the Natural Sciences and Engineering Research Council of Canada (RGPIN-2020-07212) and by the Leslie Dan Faculty of Pharmacy.

## Data Availability

MetaboMiNR can be accessed at https://cumminslab.shinyapps.io/MetaboMiNR/. The dataset used in this paper can be found in the PRIDE repository with the identifier PXD047662.

