## Supplemental User Guide for "MetaboMiNR: a web application to analyze label free quantitative proteomic experiments with a focus on metabolism and nuclear receptors"

### MetaboMiNR User Guide

**Table of Contents**

Step 1: Data upload…………………………………………………………………....2

Step 2: Global Analysis………………………………………………………………. 4

Step 3: Metabolism Miner……………………………………………………………..6

Step 4: Nuclear Receptor Miner…………………………………………………….…8

Step 5: Individual Plotter……………………………………………………………...10

### Step 1: Data upload

1. MetaboMiNR can intake LFQ data from a MaxQuant analyzed experiment. Within the MaxQuant output, you will find a file named "proteinGroups.txt". The typical path for this file is ~\combined\txt\proteinGroups.txt.

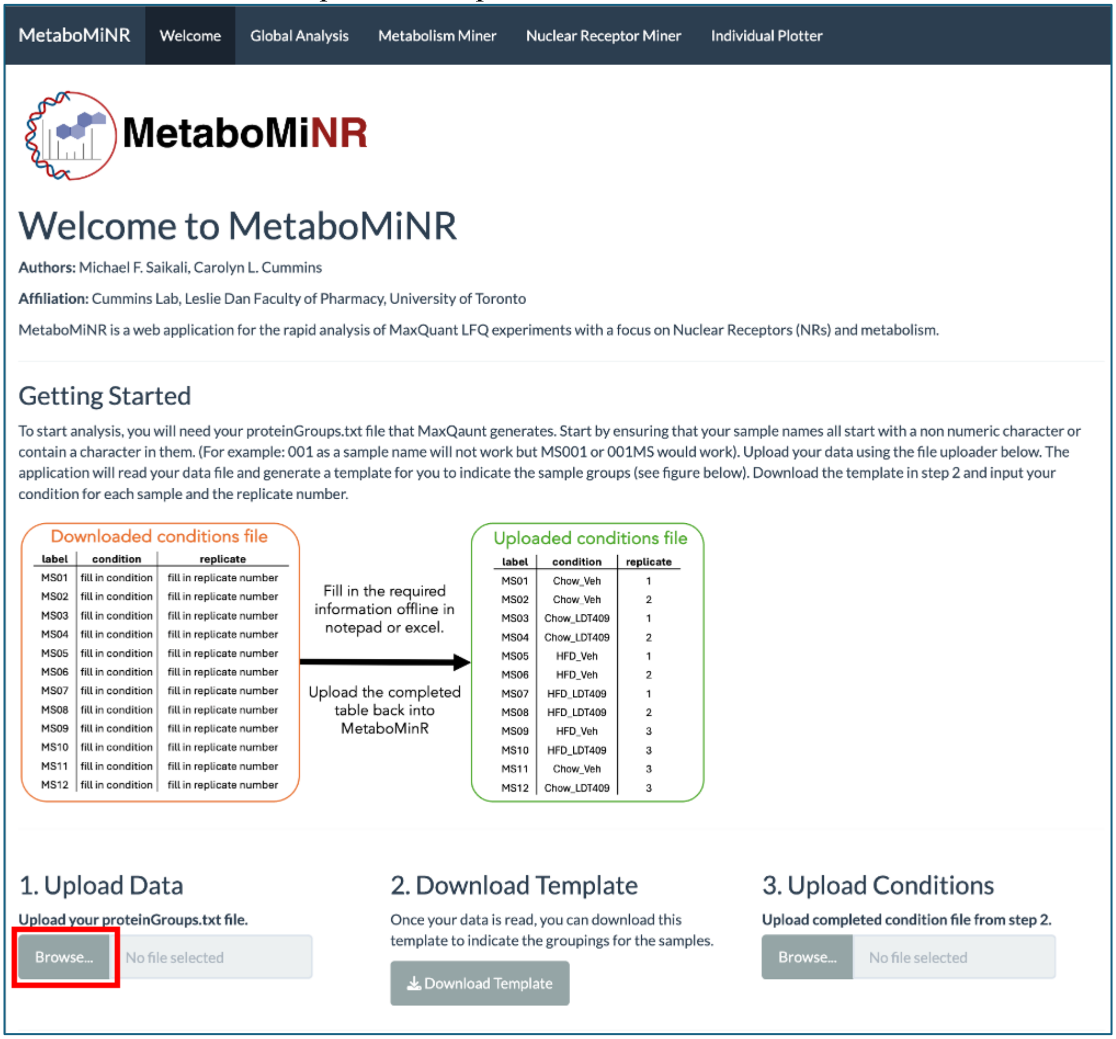


2. To upload the file, select browse under "Upload Data" and navigate to your proteinGroups.txt file. You may rename the file if you choose, but do not edit any of the columns or column names prior to upload.

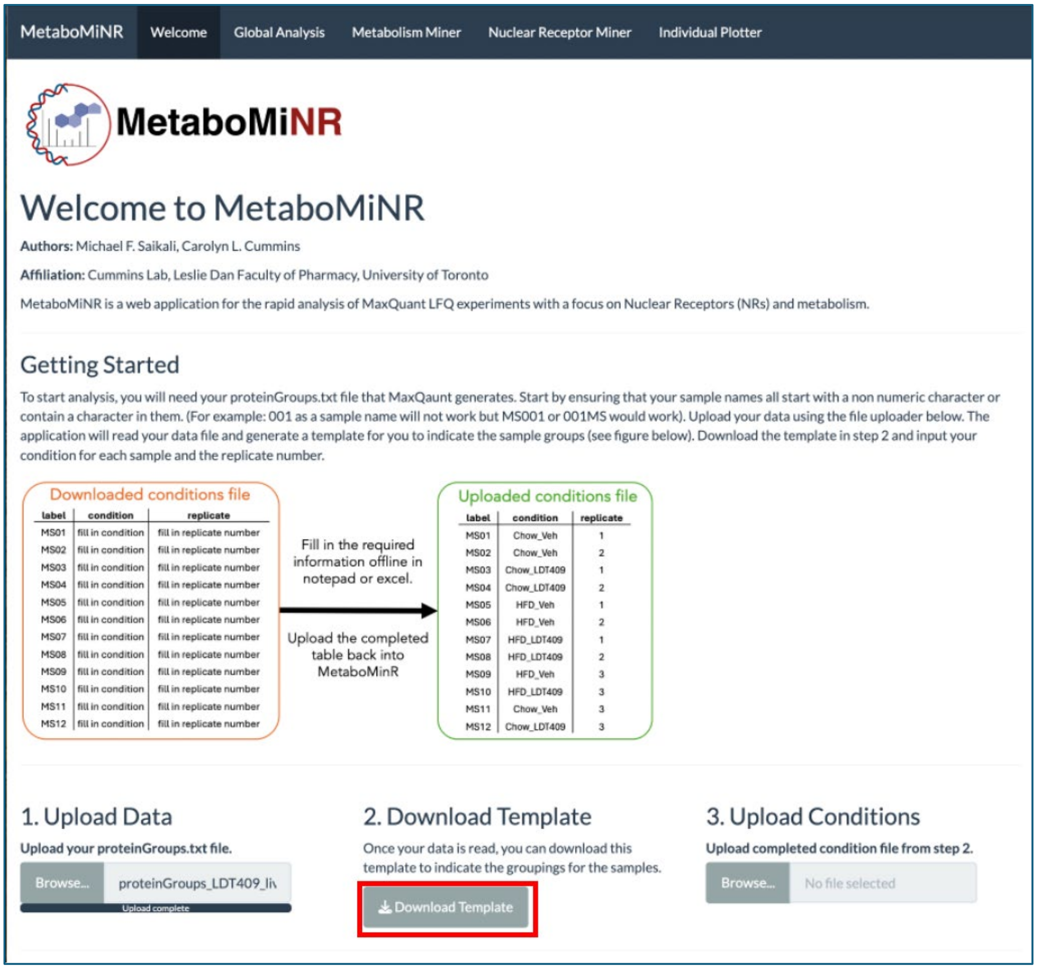


3. Once upload is complete, you will see a message under the uploader that displays “Upload complete”. You may then press “Download Template” which will automatically download a csv file named “ConditionTemplate.csv”. You may rename this file as you please.

| | A | B | C |
|---|---|---|---|
| 1 | label | condition | replicate |
| 2 | MS01 | fill in condition | fill in replicate number |
| 3 | MS02 | fill in condition | fill in replicate number |
| 4 | MS03 | fill in condition | fill in replicate number |
| 5 | MS04 | fill in condition | fill in replicate number |
| 6 | MS05 | fill in condition | fill in replicate number |
| 7 | MS06 | fill in condition | fill in replicate number |
| 8 | MS07 | fill in condition | fill in replicate number |
| 9 | MS08 | fill in condition | fill in replicate number |
| 10 | MS09 | fill in condition | fill in replicate number |
| 11 | MS10 | fill in condition | fill in replicate number |
| 12 | MS11 | fill in condition | fill in replicate number |
| 13 | MS12 | fill in condition | fill in replicate number |
| 14 | | | |

| | A | B | C |
|---|---|---|---|
| 1 | label | condition | replicate |
| 2 | MS01 | Chow_Veh | 1 |
| 3 | MS02 | Chow_Veh | 2 |
| 4 | MS03 | Chow_LDT409 | 1 |
| 5 | MS04 | Chow_LDT409 | 2 |
| 6 | MS05 | HFD_Veh | 1 |
| 7 | MS06 | HFD_Veh | 2 |
| 8 | MS07 | HFD_LDT409 | 1 |
| 9 | MS08 | HFD_LDT409 | 2 |
| 10 | MS09 | HFD_Veh | 3 |
| 11 | MS10 | HFD_LDT409 | 3 |
| 12 | MS11 | Chow_Veh | 3 |
| 13 | MS12 | Chow_LDT409 | 3 |
| 14 | | | |

4. Edit the ConditionTemplate.csv template in your editor of choice, filling in each condition avoiding spaces in the condition name. Under the replicate column, indicate for each condition which sample is replicate 1, 2, 3, etc. For each condition there cannot be duplicates of the replicate number (e.g. Chow_Veh 2, and Chow_Veh 2).

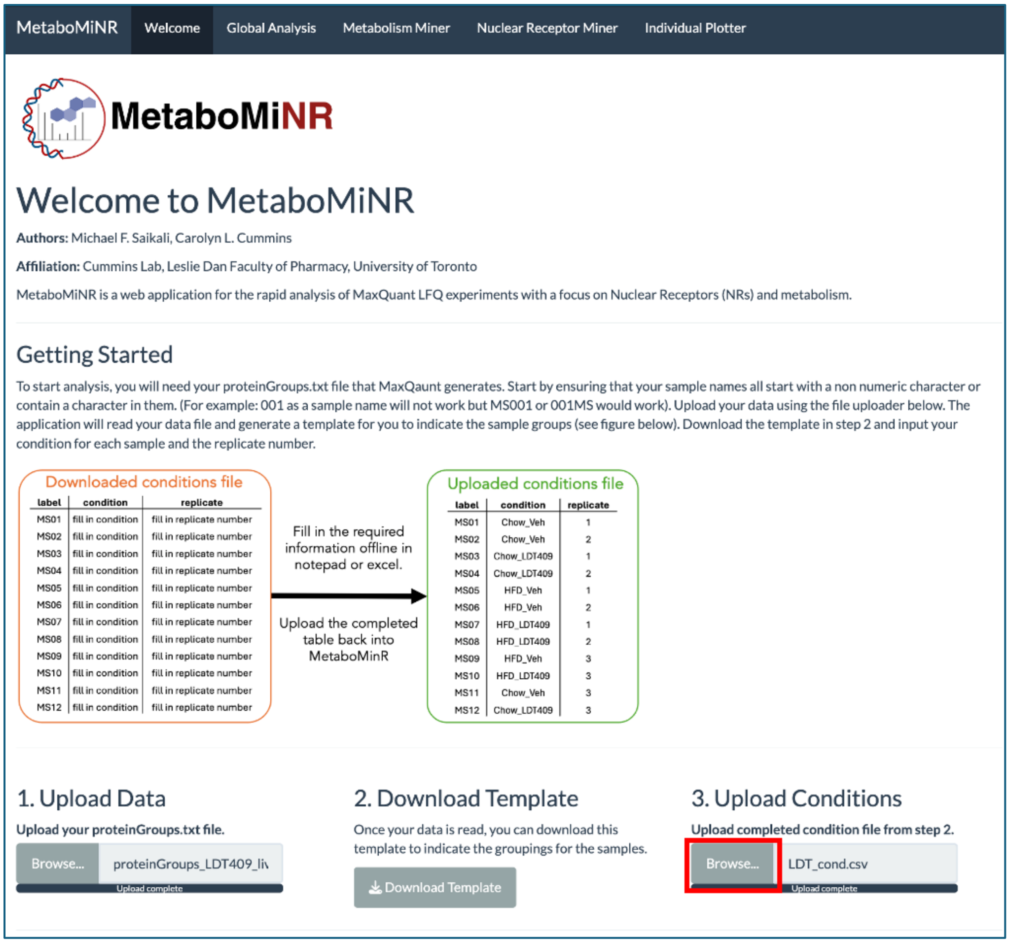


5. Upload the completed ConditionTemplate.csv file by selecting **Browse** under Upload Conditions and navigate to the completed file. Once upload is complete, you will see a progress bar that reads “Upload complete”.

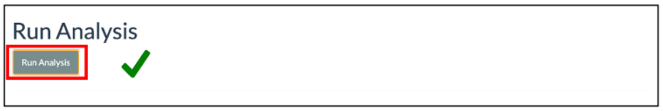


6. To begin analysis, press on **Run Analysis** and wait until a green checkmark appears next to the button. Once this green checkmark appears, you may navigate to other pages.

### Step 2: Global Analysis

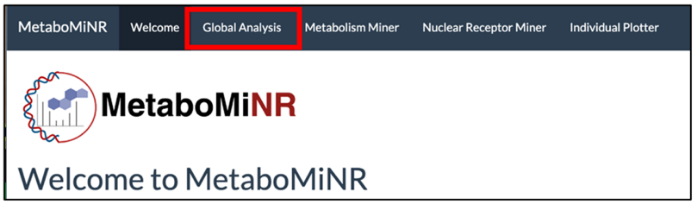


7. Using the navigation bar at the top of the page, navigate to the **Global Analysis** page.

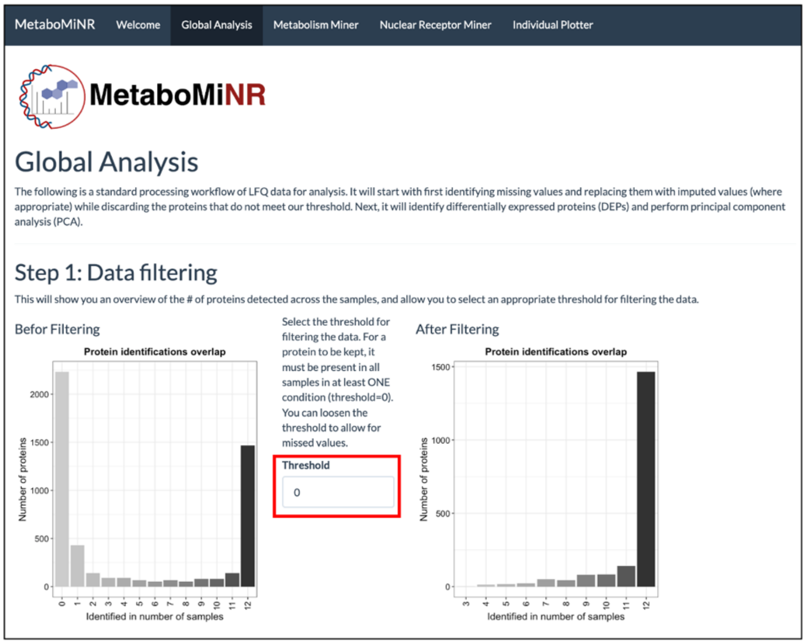


8. Under the data filtering section, you will find a plot showing the number of proteins identified in each sample before and after filtering for valid data. Between them is a threshold value that you can adjust. The default selection is 0 which means that for a protein to be kept in the dataset, it must be detected in *ALL* replicates of at least *ONE* condition. Increasing the threshold decreases that requirement by 1. For example, when in a 4-replicate experiment, threshold = 1 requires the protein to be detected in at least 3 out of 4 of the replicates in at least one condition.

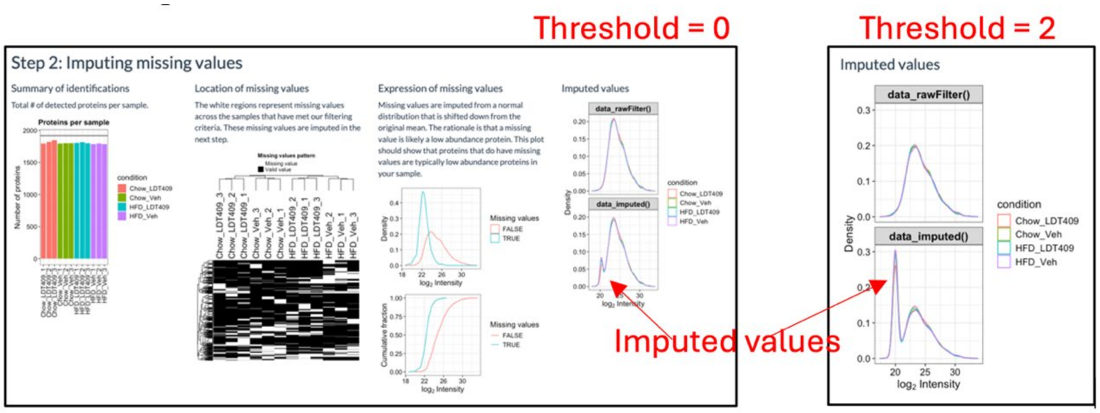


9. The next step shows the process of imputing missing values. Missing values are usually imputed in proteomic analyses to preserve the ability to perform statistical analyses on proteins that are not present in all conditions. These values are imputed as a random value from a normal distribution taken 1.8 standard deviations below the mean of the observed values. The heatmap displays where the missing values are, allowing the user to spot a poor sample if it is consistently missing values. The histograms on the right labeled data_rawFilter() and data_imputed() show the distribution of intensities before and after imputing, respectively. You will see a new small distribution or shoulder appear on the left of the histogram representing the imputed values. The figure above to the left shows the histogram for imputing threshold of 0 compared to an imputing threshold of 2 on the right. Increasing the threshold increases the number of imputed values. ***The imputed value peak height should not exceed the real values.*** Thus, in the image above, the threshold of 2 is too high.

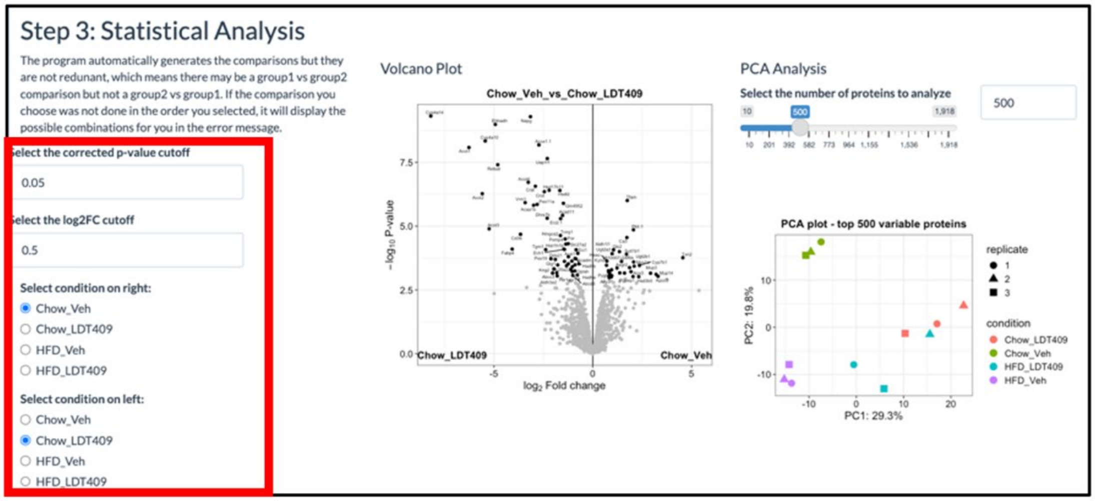


10. The next step includes the statistical analysis. The default cutoff for the adjusted p-value is 0.05, however you may decrease this to make the cutoff more stringent. The $log_2FC$ cutoff is set to 0.5 by default meaning it will only highlight proteins that were greater than 0.5 or less than -0.5.

Valid contrasts are: 'Chow_LDT409_vs_HFD_LDT409', 'Chow_LDT409_vs_HFD_Veh', 'Chow_Veh_vs_Chow_LDT409', 'Chow_Veh_vs_HFD_LDT409', 'Chow_Veh_vs_HFD_Veh', 'HFD_Veh_vs_HFD_LDT409'

11. You may change the conditions plotted by selecting a different condition using the radio buttons. While the application does every possible comparison, it only does it once (e.g. it will do Chow_Veh vs Chow_LDT409 but not Chow_LDT409 vs Chow_Veh). If you have a selected an invalid comparison, reverse the selection or check the list of valid contrasts that appear with the error message.

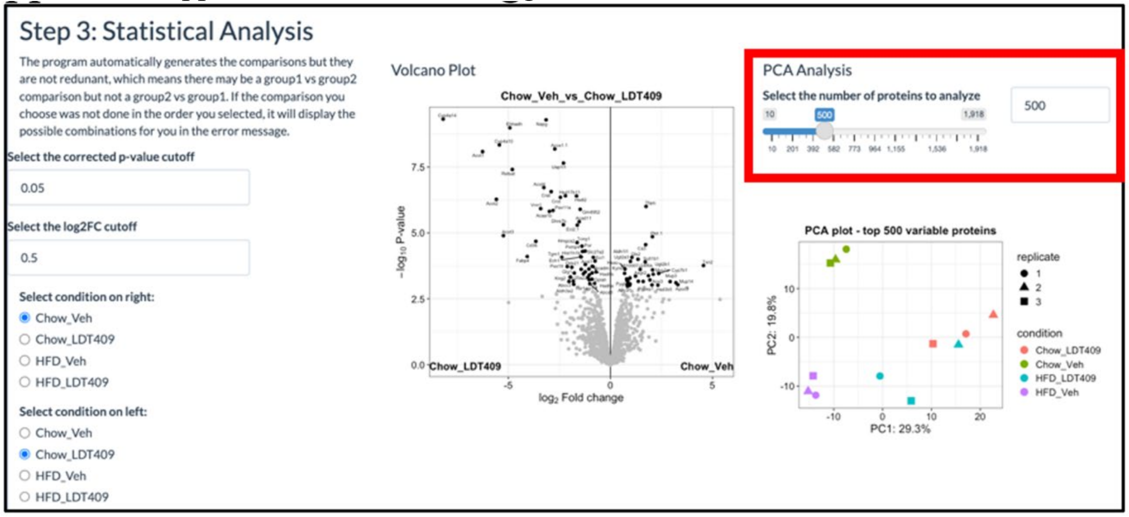


12. The default setting for the PCA plot is to generate it using only the top 500 most variable proteins. You may use the slider bar or the text input to increase this to a maximum of the number of proteins detected in your samples allowing you to produce a PCA plot of the entire dataset.

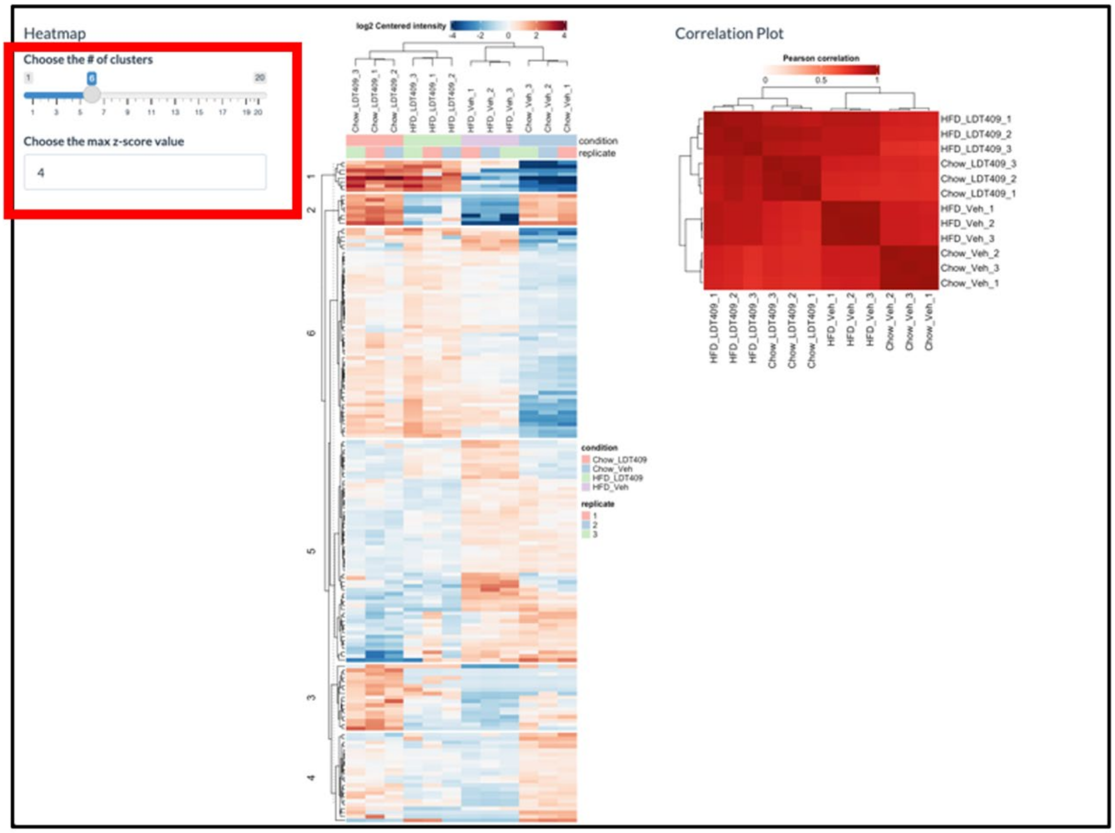

13. The next step produces a k-means clustered heatmap. The default selection for the number of clusters is 6, but you may use the slider to adjust the number of clusters and the heatmap will adjust in real time. Additionally, you may adjust the color scaling by increasing or decreasing the maximum z-score value which sets the limits for the legend.

### Step 3: Metabolism Miner

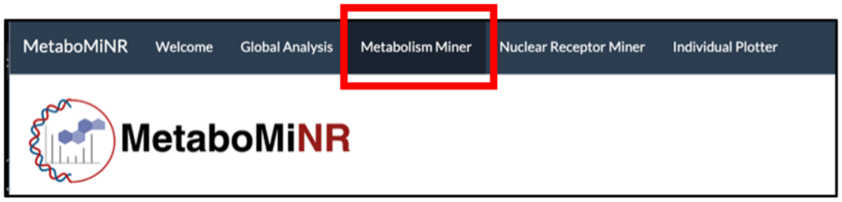


14. Navigate to the Metabolism Miner tool using the navigation bar.

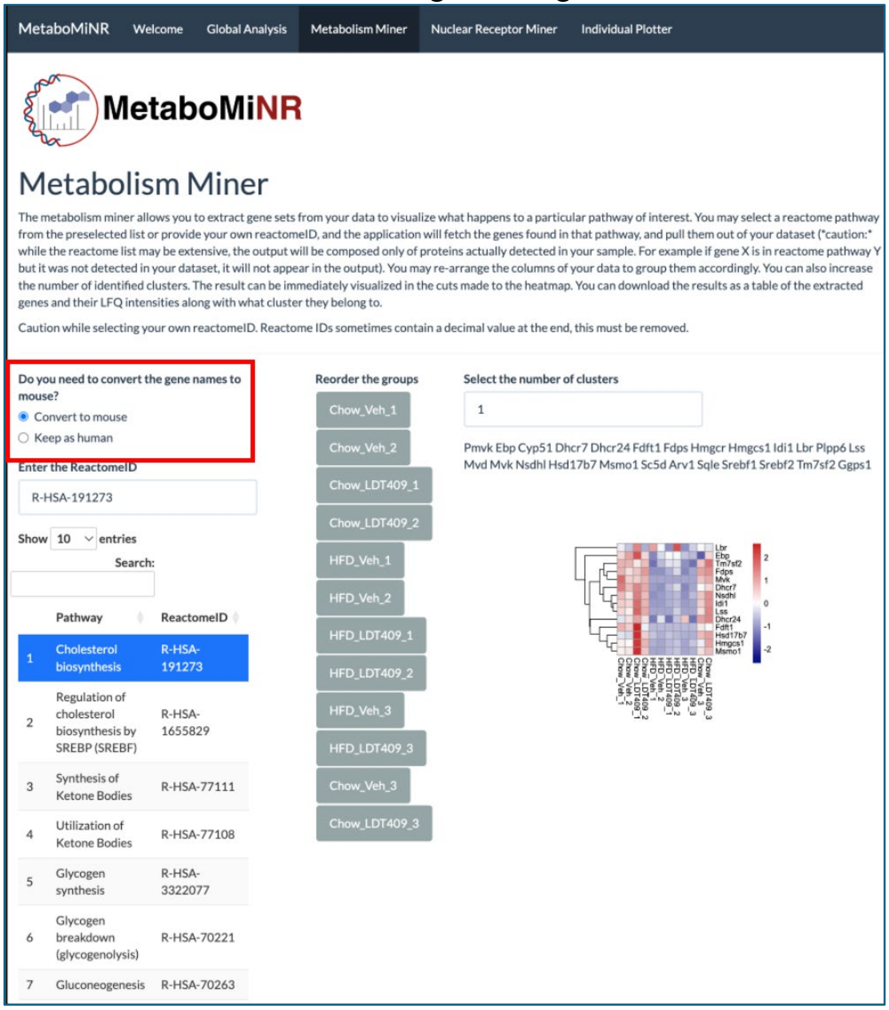


15. The Metabolism Miner pulls genes from Reactome which are stored as human gene names. The default setting in MetaboMiNR is to convert these gene names to their mouse orthologues. If you are dealing with a human sample, change this to “Keep as human”.

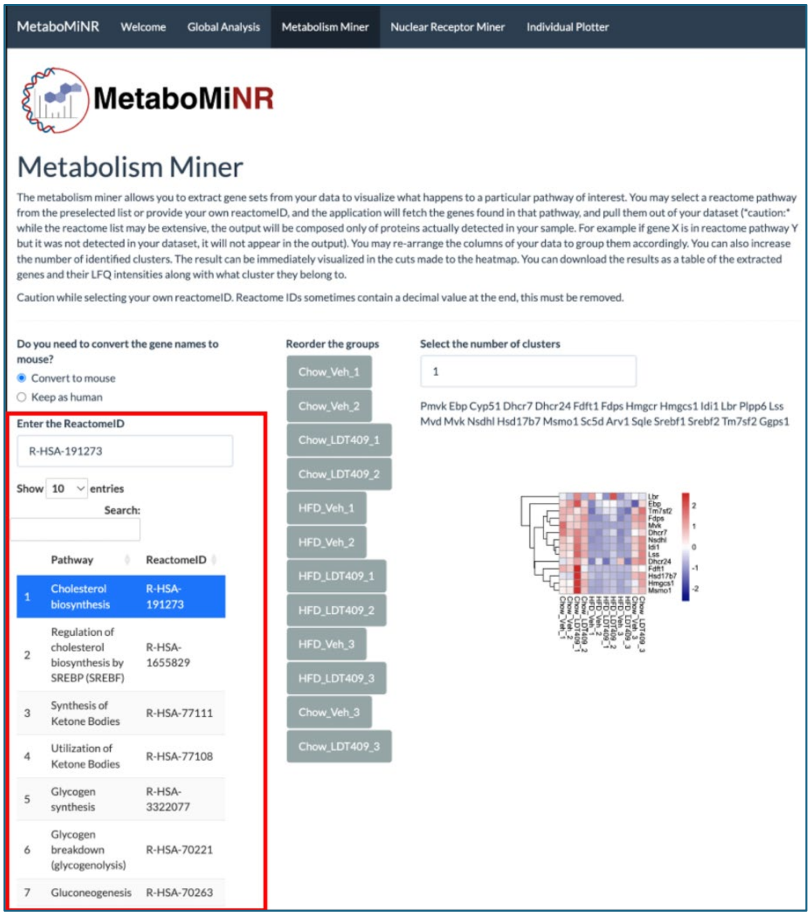


16. To select a pathway, you can choose from the preselected pathways or enter your own ReactomeID in the textbox. Some ReactomeIDs will have a decimal and numbers at the end, remove these prior to inputting the ReactomeID into the textbox.

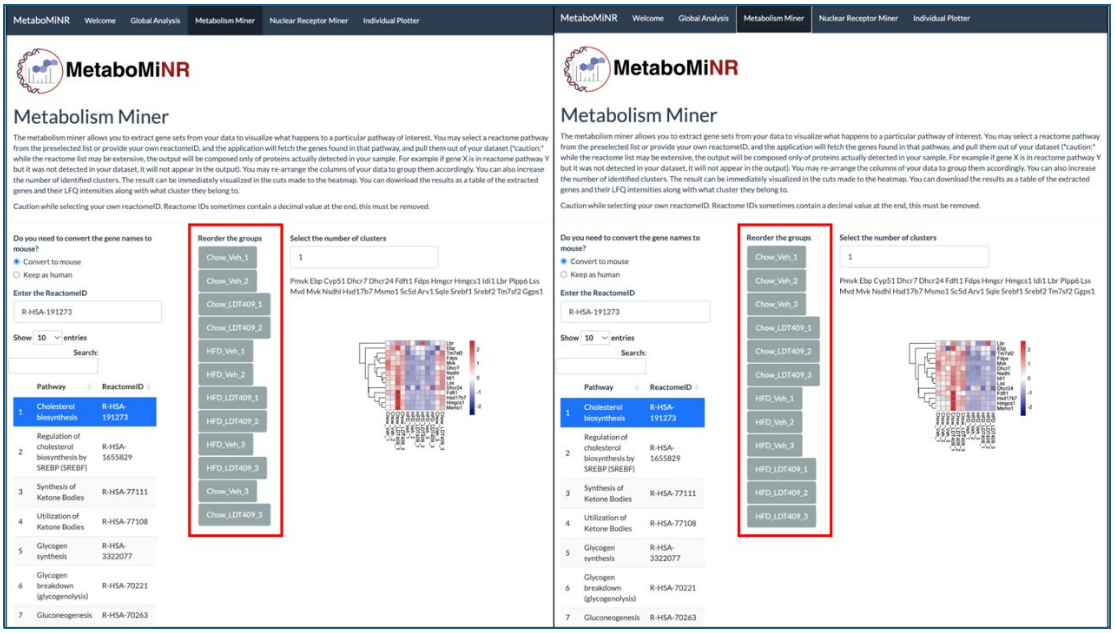

17. To re-arrange the heatmap, simply drag and drop the sample name tiles into the desired order and the heatmap will re-arrange.

Download Results

Search:

| | Cluster | Chow_Veh_1 | Chow_Veh_2 | Chow_Veh_3 | Chow_LDT409_1 | Chow_LDT409_2 | Chow_LDT409_3 | HFD_Veh_1 | HFD_Veh_2 | H |
|---|---|---|---|---|---|---|---|---|---|---|
| Lbr | 1 | 2216200 | 1796900 | 2278000 | 3798600 | 1549127 | 2147700 | 3458400 | 2296800 | |
| Ebp | 1 | 15585000 | 18293000 | 6494100 | 22087000 | 15502000 | 15317000 | 8552700 | 11781000 | |
| Tm7sf2 | 1 | 11883000 | 11763000 | 10554000 | 8939400 | 11201000 | 14777000 | 1301329 | 4707700 | |
| Fdps | 1 | 256480000 | 196970000 | 275770000 | 225720000 | 292280000 | 303250000 | 25422000 | 20024000 | |
| Mvk | 1 | 13552000 | 8146300 | 8665200 | 8129400 | 9843700 | 10318000 | 1301329 | 1226478 | |
| Dhcr7 | 1 | 15314000 | 11821000 | 8141000 | 14949000 | 12785000 | 10313000 | 2897200 | 2853200 | |
| Nsdhl | 1 | 61771000 | 43985000 | 59044000 | 86296000 | 63528000 | 79192000 | 1301329 | 5147600 | |
| Idi1 | 1 | 68499000 | 66318000 | 68941000 | 119630000 | 81952000 | 100780000 | 1301329 | 1226478 | |
| Lss | 1 | 12068000 | 9759800 | 11450000 | 22611000 | 15816000 | 19097000 | 1301329 | 1226478 | |
| Dhcr24 | 1 | 4035700 | 5781000 | 7532500 | 7505900 | 5787500 | 8525100 | 1301329 | 4201800 | |

Showing 1 to 10 of 14 entries — Previous 1 2 Next

18. At the bottom of the page you will find a table with a download button that allows you to download the extracted data for downstream analysis.

### Step 4: Nuclear Receptor Miner

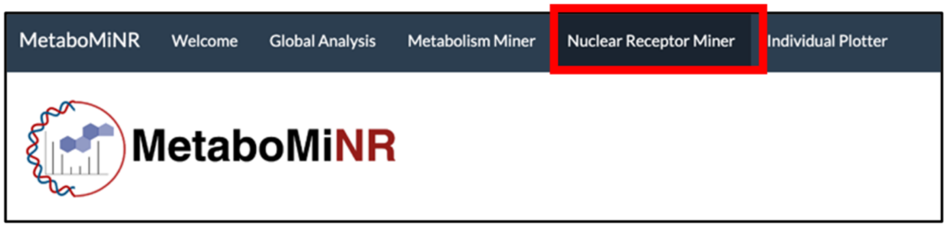


19. Navigate to the Nuclear Receptor Miner page.

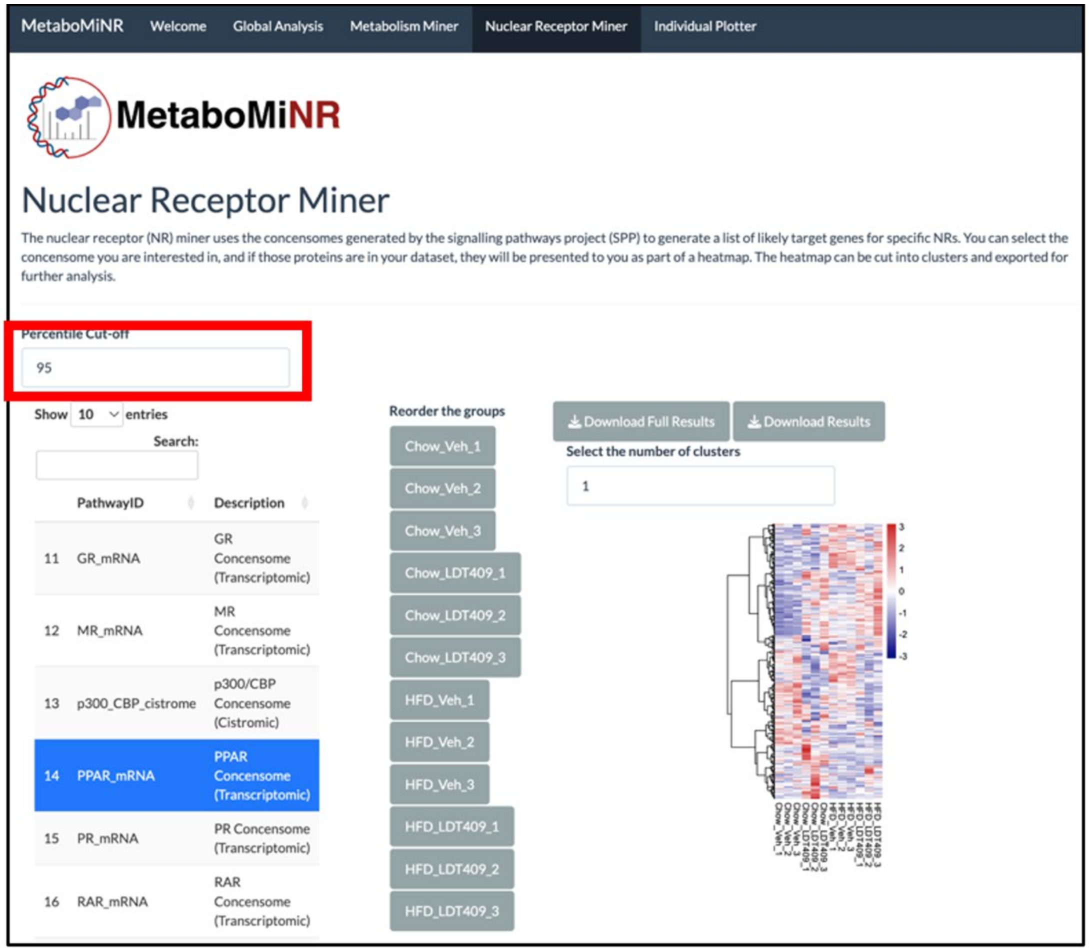


20. Similar to the Metabolism Miner page, you may select a pathway to extract and a heatmap will be produced. The order of the columns can be adjusted by dragging the sample name tiles around. The default percentile cutoff is 95% meaning only the 95$^{th}$ percentile and above is taken as a target gene in that consensome. You can make this more stringent by increasing the value.

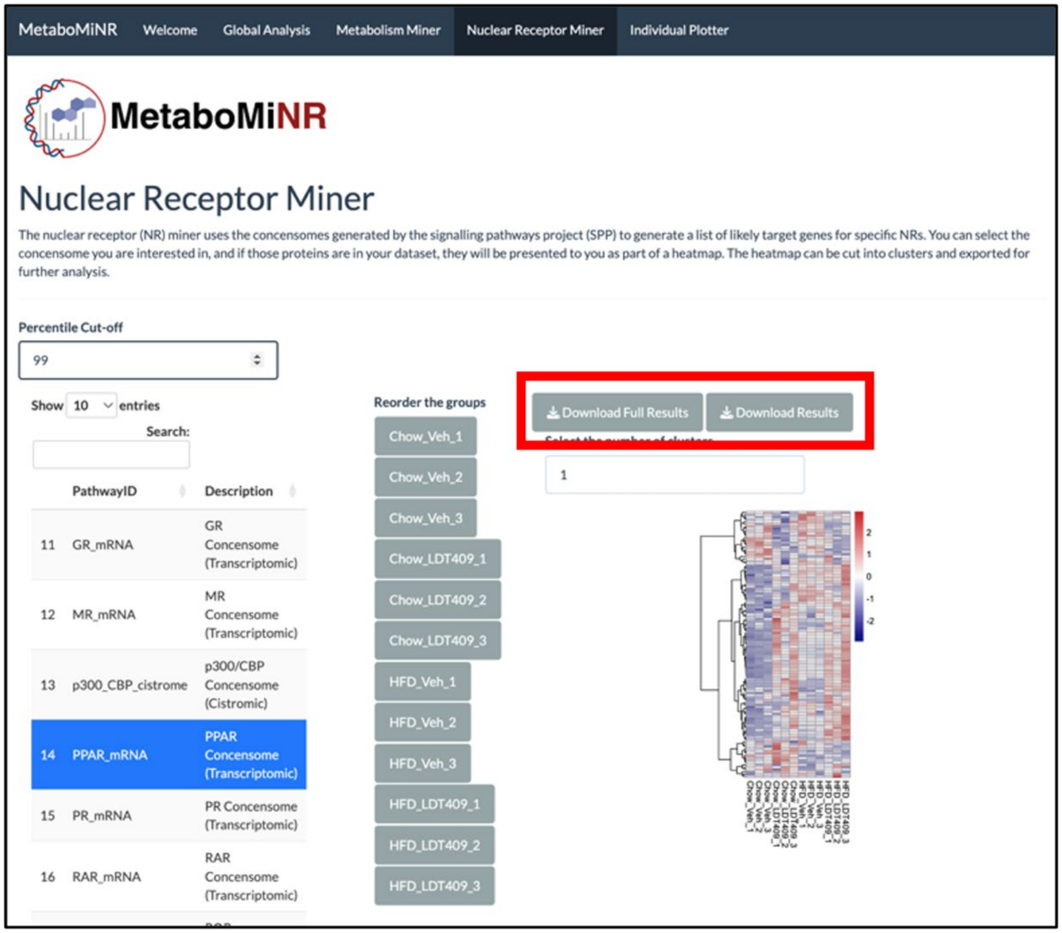


21. By selecting Download Results, you will download a table of the data shown in the heatmap. By selecting Download Full Results, you will download a table containing the entire dataset, with the adjusted p-values for each comparison, and appended to it will be a table of the consensome results. See the figure below.

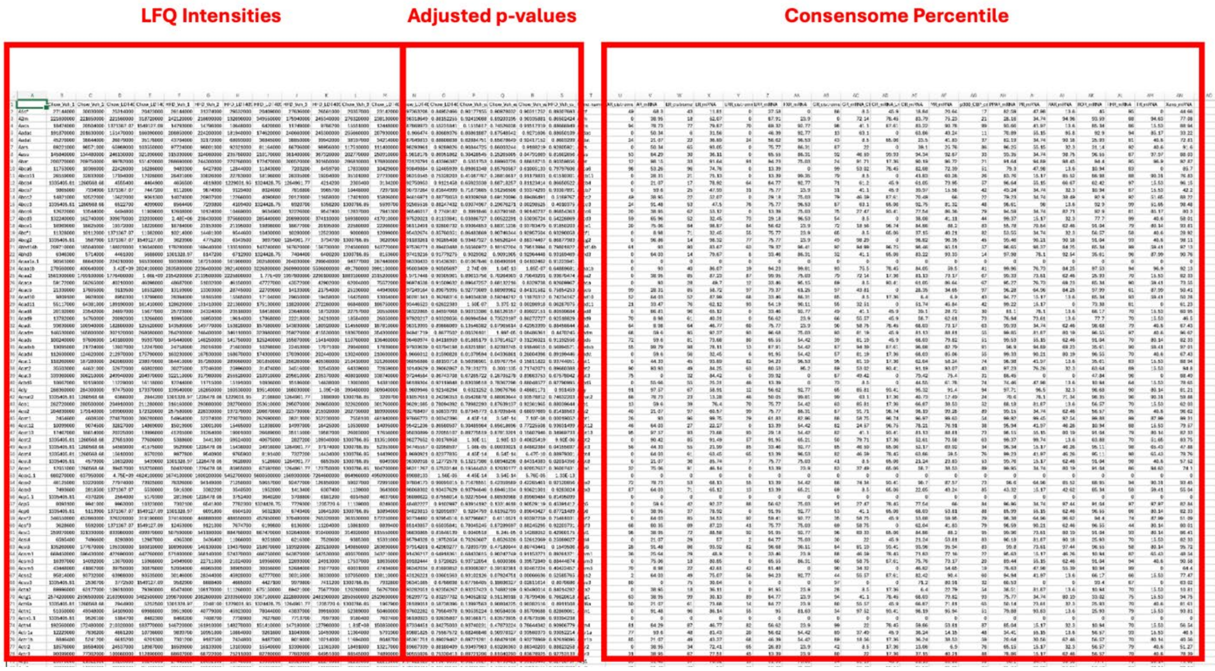


22. Offline, you can use this table to filter for only significantly altered proteins and plot their consensomes results.

### Step 5: Individual Plotter

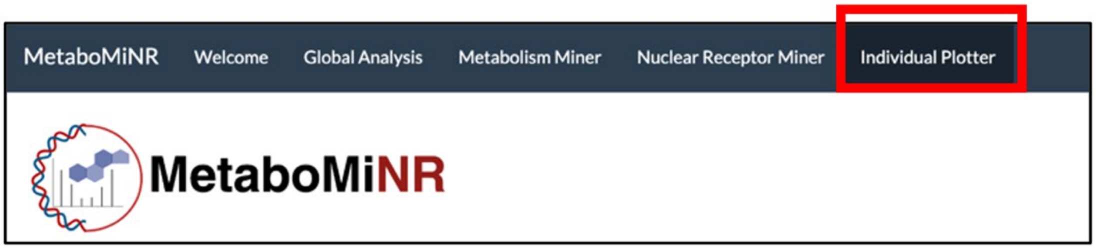


23. Navigate to the final tab, the Individual Plotter.

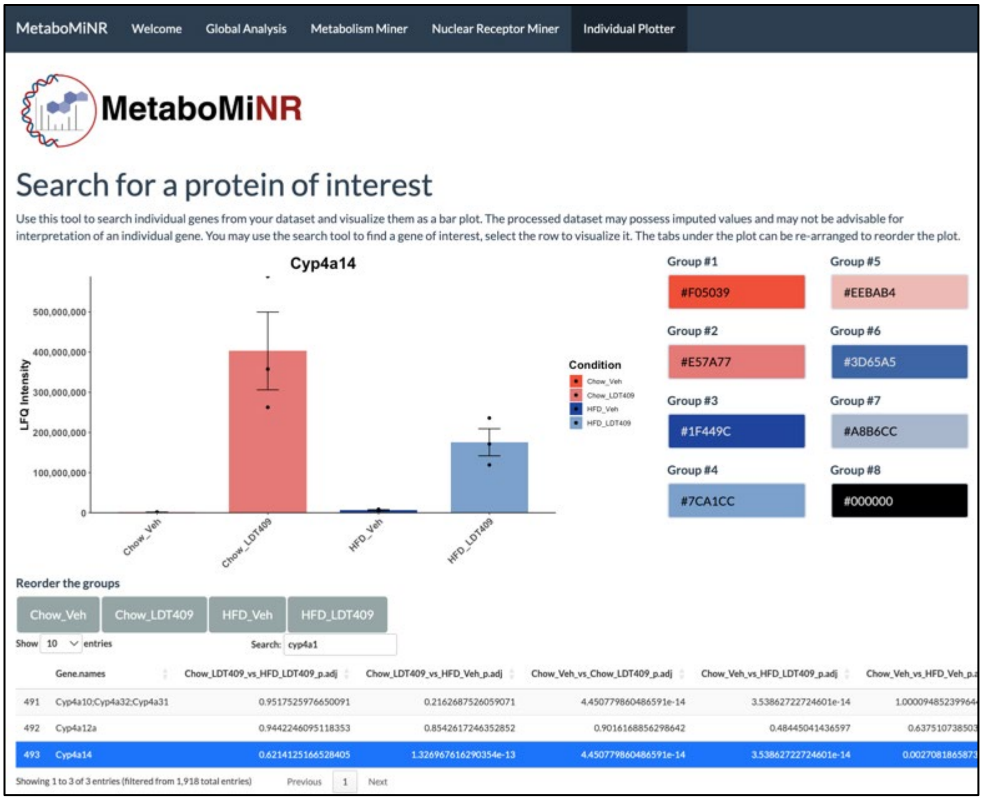


24. Use the search bar to locate a gene. The table will automatically update as you narrow the search. To plot the gene, select the row and it will be highlighted in blue and automatically plotted. The p-values cannot be plotted directly on the figure but can be found in the table where you selected the gene.

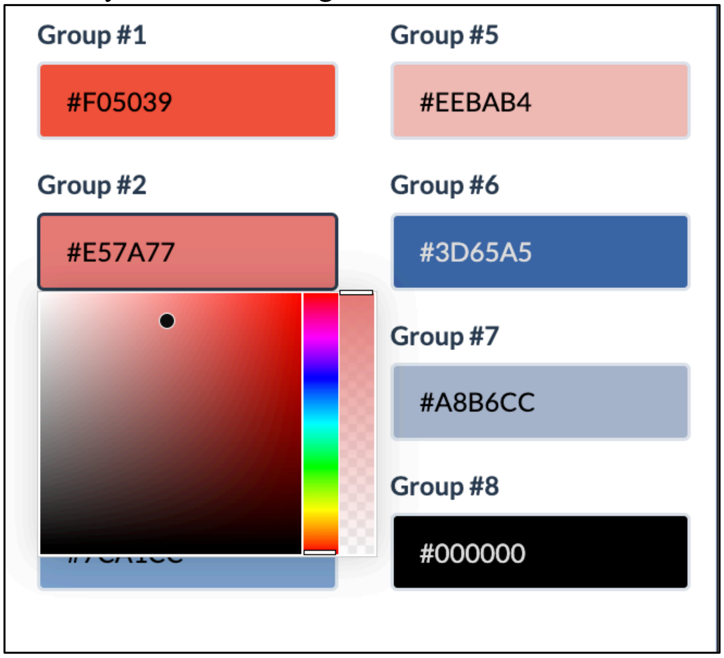


25. To adjust the color of the plot you can select the group you want to edit and use the color picker to change the color and opacity.

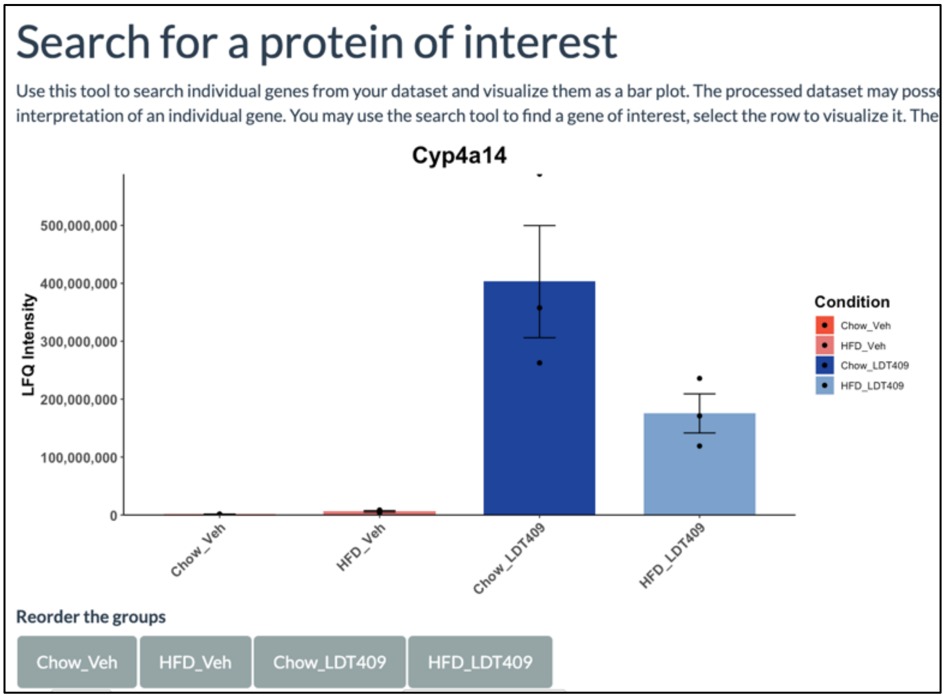


26. Finally, to adjust the order of the bars, drag the grey tiles around into the order that you would like them to appear.
